# Optimal information transfer in enzymatic networks: A field theoretic formulation

**DOI:** 10.1101/130112

**Authors:** Himadri S. Samanta, Michael Hinczewski, D. Thirumalai

## Abstract

Signaling in enzymatic networks is typically triggered by environmental fluctuations, resulting in a series of stochastic chemical reactions, leading to corruption of the signal by noise. For example, information flow is initiated by binding of extracellular ligands to receptors, which is transmitted through a cascade involving kinase-phosphatase stochastic chemical reactions. For a class of such networks, we develop a general field-theoretic approach in order to calculate the error in signal transmission as a function of an appropriate control variable. Application of the theory to a simple push-pull network, a module in the kinase-phosphatase cascade, recovers the exact results for error in signal transmission previously obtained using umbral calculus (Phys. Rev. X., **4**, 041017 (2014)). We illustrate the generality of the theory by studying the minimal errors in noise reduction in a reaction cascade with two connected push-pull modules. Such a cascade behaves as an effective three-species network with a pseudo intermediate. In this case, optimal information transfer, resulting in the smallest square of the error between the input and output, occurs with a time delay, which is given by the inverse of the decay rate of the pseudo intermediate. Surprisingly, in these examples the minimum error computed using simulations that take non-linearities and discrete nature of molecules into account coincides with the predictions of a linear theory. In contrast, there are substantial deviations between simulations and predictions of the linear theory in error in signal propagation in an enzymatic push-pull network for a certain range of parameters. Inclusion of second order perturbative corrections shows that differences between simulations and theoretical predictions are minimized. Our study establishes that a field theoretic formulation of stochastic biological signaling offers a systematic way to understand error propagation in networks of arbitrary complexity.

## INTRODUCTION

Cell signaling involves the ability of cells to detect changes in the environment and respond to them [1–6], a fundamental necessity of living systems. Several signaling networks involve proteins, which switch between active and inactive states. By quantitatively describing how different signaling proteins are functionally linked, we can understand the behavior of signaling pathways, and the associated bandwidth that determines fidelity of information transfer [7]. In typical enzymatic networks, environmental information is transmitted into the cell interior through cascades of stochastic biochemical reactions [8]. Noise inevitably propagates through the cascade, potentially corrupting the signal. Depending on the parameters, small changes in the input can be translated into large (but noise corrupted) output variations. The amplification is essential but it must also preserve the signal content to be useful for downstream processes. The signaling circuit, despite operating in a noisy environment, needs to maintain high fidelity between output and the amplified input [9]. Over the years concepts in information theory have been adopted to assess the fidelity of signal transmission in the context of biochemical network [10, 11]. Several studies have used mutual information between input and output signals to quantify the reliability of signal transduction [12–17]. The formalism has been applied to the study of a variety of networks including cascades and networks with feedback [12]. These and other studies have expanded over understanding of the fidelity of information transfer in biological networks in which both noise and copy number fluctuations are important.

In a recent paper [18], we considered the problem of how to extract information faithfully from noisy signals using mathematical methods developed in the context of communication theory developed over sixty years ago by Wiener [19] and independently by Kolmogorov [20]. The Wiener-Kolmogorov (WK) approach has since proven a useful tool in a variety of contexts in biological signaling [9, 21, 22]. The WK theory, reformulated by Bode and Shannon [11], assumes that the input and output are continuous variables that describe stationary stochastic processes. The goal of approach is to minimize the mean squared error between the input and output signals, but the optimization is restricted to the space of only linear noise filters. Recently, we developed an analytic formalism of general validity to overcome some of the limitations of the WK theory based on exact techniques involving umbral calculus [23]. We illustrated the efficacy of the non-linear theory with applications to push-pull network and its variants including instances when the input is time-dependent.

The use of non-standard mathematics in the form of umbral calculus, perhaps, obscures the physics of optimal filtering in biological networks in which the effects of non-linearities in signal amplification have to be considered. Here, we develop an alternate general formalism based on a many body formulation of reaction diffusion equations introduced by Doi and Peliti [24, 25]. This formulation converts the signal optimization problem to a standard field theory, allowing us to calculate the response and correlation functions by standard methods. The advantage of this formalism is that both discrete and continuum cases can be studied easily. Non-linear contributions can be obtained using systematic diagrammatic perturbation scheme for an arbitrary network. Networks where temporal dynamics are coupled with spatial gradients in signaling activities, which regulate intracellular processes and signal propagation across the cell, can also be investigated using the present formalism. Application of the theory to a push-pull network and a simplified biochemical network recovers the exact results obtained in our previous study. We also extend the formalism to solve signal transduction in a cascade, which serves as a model for a variety of biological networks. The formalism is general and is applicable to arbitrary networks with feedback, time delay and special variations [26]. Our work exploits standard methods in physics, illustrating the usefulness of a field theoretic formulation at the interface of communication theory and biology.

## THEORY

### Linear Push-Pull Network

In order to develop the many body formalism for a general signaling network, we first consider a simple model. The concepts and the general diagrammatic expansion developed in this context, lays the foundation for applications to more complicated enzymatic networks as well as signaling cascades. In a typical signaling pathway, for example the mitogen activated protein kinase (MAPK) [27, 28] pathway, external and environmental fluctuations activate a cascade of enzymatic reactions, thus transmitting information across the membrane in a sequential manner. Each step involves activation of kinases by phosphorylation reaction and deactivation by phosphatases [29–33]. A truncated version of such a cascade is a single step (Fig.1), which we refer to as a push-pull network [18]. In this signaling network, there are only two chemical species. One is *I*(*t*) (the ”input”) and the other is *O*(*t*) (the ”output”) whose production depends on *I*(*t*). The upstream pathway, which serves as an external signal, creates the species *I* by the re-action 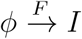 with an effective production rate *F*. The output *O* is a result of the reaction 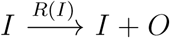, with a rate *R*(*I*(*t*)) that depends on the input. The species are deactivated through 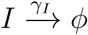 and 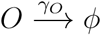 with rates *γ*_*I*_ and *γ*_*O*_ respectively, mimicking the role of phosphatases (Fig.1). The input varies over a characteristic time scale 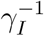, fluctuating around the mean value 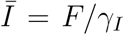. The degradation rate sets the time scale 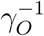 over which *O*(*t*) responds to changes in the input.

**FIG. 1:**
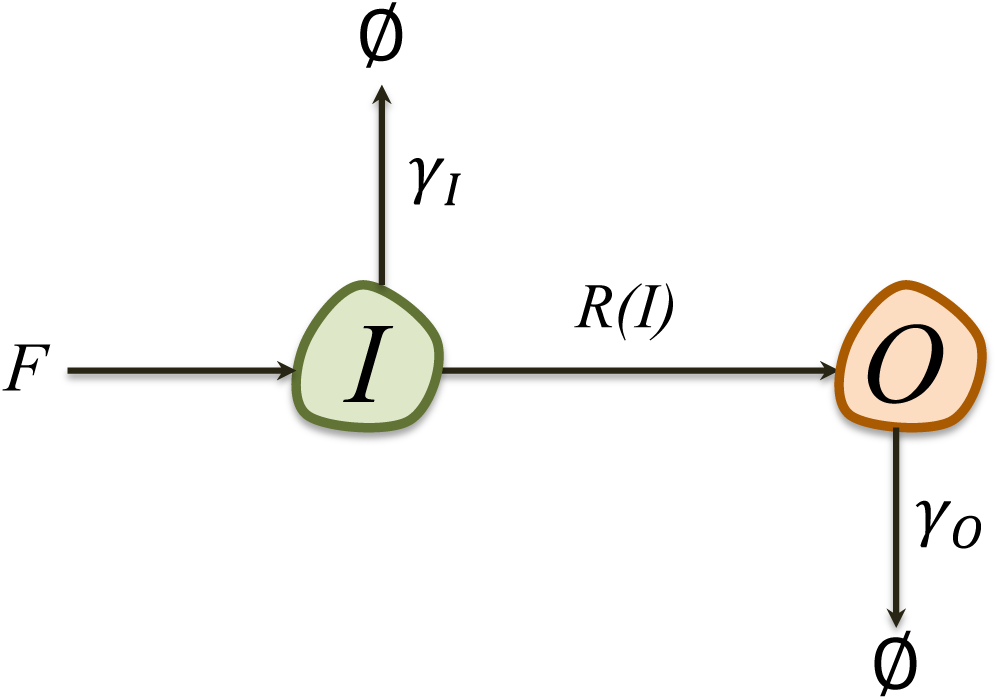
Schematic of a push-pull network, involving an input species *I* and output species *O*. The production of *O* from *I* is controlled by the rate function *R*(*I*). The degradation rates for *I* and *O* are *γ*_*I*_ and *γ*_*O*_, respectively.

The chemical Langevin equations describing the changes in *I* and *O* are,

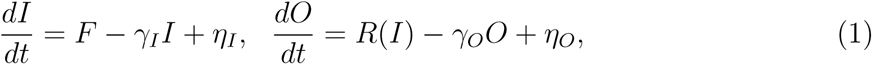

where *η*_*I*_ and *η*_*O*_ are Gaussian white noise with zero mean (⟨*η*_*α*_⟩ = 0) and correlation 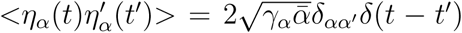 with *α* = *I, O* and 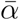 is the mean population *α*. For small fluctuations, 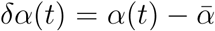, Eq.(1) can be solved using a linear approximation for the rate function 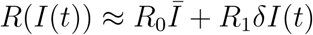, with coefficients *R*_0_, *R*_1_*>*0. The result is

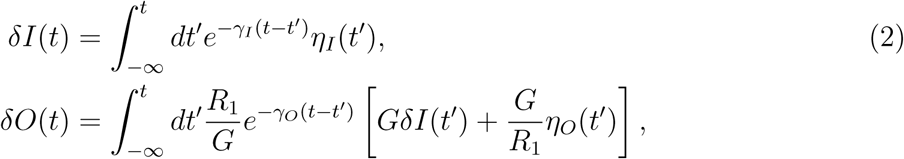

where in the second line an arbitrary scaling factor *G* has been introduced. The solution for *δO*(*t*) has the structure of a linear noise filter equation; 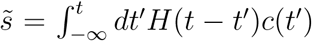, with *c*(*t*) = *s*(*t*) + *n*(*t*). The signal *s*(*t*) = *GδI*(*t*) together with the noise term 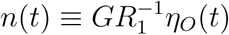 constitute the corrupted signal, *c*(*t*). The output 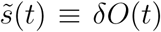 is produced by convolving *c*(*t*) with a linear kernel 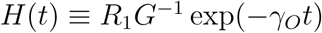, which filters the noise. As a consequence of causality, the filtered output 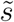 at time *t* depends only on *c*(*t′*) from the past.

The primary goal in transmitting signal with high fidelity is to devise an optimal causal filter, *H*_*opt*_(*t*), which renders 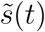 as close to *s*(*t*) as possible. In a remarkable development, Weiner [19] and Kolmogorov [20] independently discovered a solution to this problem in the context of communication theory, which launched the modern era in signal decoding from time series. In particular, WK proposed a solution that minimizes the square of the differences between 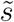 and *s*(*t*) by seeking an optimal filter *H*_*W*__*K*_(*t*) among all possible linear filters. In the push-pull network, this means having *δO*(*t*) reproduce as accurately as possible the scaled input signal *GδI*(*t*). For a particular *δI*(*t*) and *δO*(*t*), the value of the mean squared error 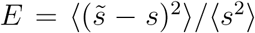 is smallest when 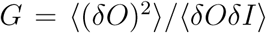, which we identify as a gain factor. In this case, 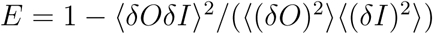.

The optimal causal filter *H*_*W*__*K*_ satisfies the following Wiener-Hopf equation[9, 18],

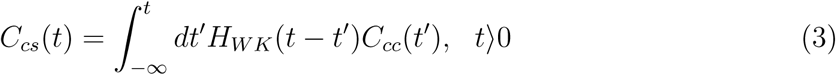

where *C*_*xy*_(*t*) = ⟨*x*(*t′*)*y*(*t′* + *t*)⟩ is the correlation between points in the time series *x* and *y*, assumed to depend only on the time difference *t − t′*. We can evaluate the correlation functions *C*_*cs*_ and *C*_*cc*_ using Eq. (2), and substituting these solutions in Eq. (3), the optimal filter function can be solved by assuming a generic ansatz, 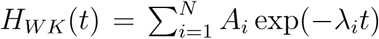. The unknown coefficients, *A*_*i*_, and the associated rate constants *λ*_*i*_ are found by comparing the left and right hand sides of Eq. (3). Elsewhere [18], we showed that 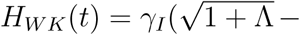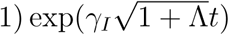. The conditions for achieving WK optimality, *H*(*t*) = *H*_*WK*_(*t*), are [18],

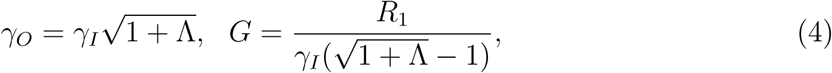

leading to the minimum relative error,

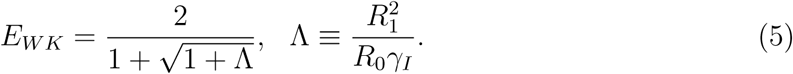

The fidelity between the output and input is described through a single dimensionless optimality control parameter, Λ, which can be written as Λ ≡ (*R*_0_*/γ*_*I*_)(*R*_1_*/R*_*o*_)^2^. The first term, *R*_0_*/γ*_*I*_, is a burst factor, measuring the mean number of output molecules produced per input molecule during the active lifetime of the input molecule. The second term, (*R*_1_*/R*_*o*_)^2^, is a sensitivity factor, reflecting the local response of the production function *R*(*I*) near 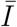 (controlled by the slope 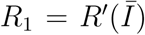) relative to the production rate per input molecule 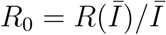.

In our recent work [18], we extended the WK approach to include non-linearity and the discrete nature of the input and output molecules *I* and *O* [18]. Both these considerations are relevant in biological circuits where *R*(*I*) is non-linear and the copy numbers of *I* and *O* are likely to be small. Starting from the exact master equation, valid for discrete populations and arbitrary *R*(*I*), we rigorously solved the original optimization problem for the error *E* between output and input using the principles of umbral calculus[23]. The main results are as follows. For any arbitrary function expanded as,

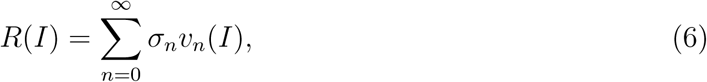

with 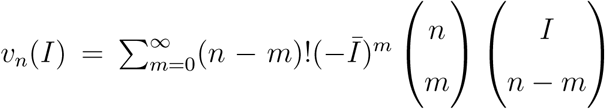and 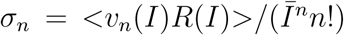, the relative error can be expressed by an exact expression,

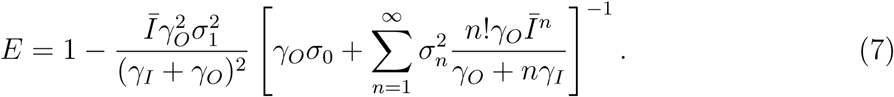

The expression above is bounded from below by

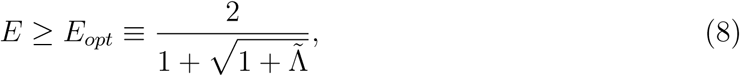

where 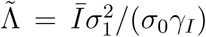. The equality is only reached when 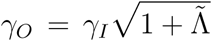 and *R*(*I*) is an optimal linear filter of the form, 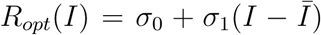, with all *σ*_*n*_ = 0 for *n ≥* 2. Obtaining the lower bound is important for noise reduction in biological networks as it provides insights into energy costs required to reduce the error [10].

### Field theoretic formulation

In order to generalize the results in our previous study [18] to arbitrary regulatory networks, we adopt a many body approach pioneered by Doi and Peliti[24, 25]. Such an approach has been used in the study of a variety of reaction diffusion equations [34, 35]. Besides suggesting plausible new ways of examining how signals are transmitted in biochemical reaction networks, the current theory shows how standard field theoretic methods can be adopted for use in control theory. By way of demonstrating its utility, we rederive the exact analytical solution (Eq.(7)) for the relative error in the push-pull network. In the Doi-Peliti formalism, the configurations at time *t* in a locally interacting many body system are specified by the occupation numbers of each species on a lattice site *i*. In our case, *I*_*i*_ is the input population and *O*_*i*_ is the output population. As a consequences of the stochastic dynamics, the on-site occupation numbers are modified. Arbitrarily many particles of either population are allowed to occupy any lattice site. In other words, *I*_*i*_, *O*_*i*_ = 0, 1, *… ∞*. The master equation for the local reaction scheme that governs the time evolution of the configurational probability with *I*_*i*_ input and *O*_*i*_ output at site *i* at time *t* is obtained through the balance of gain and loss terms. The result is,

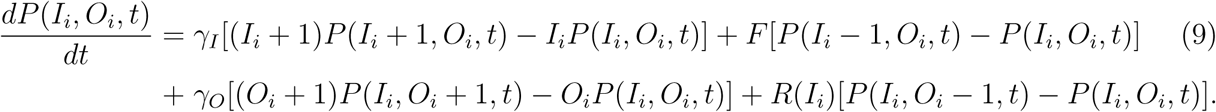

We use the Fock space representation to account for the changes in the site occupation number by integer values for the chemical reactions describing the network. Following Doi and Peliti, we introduce the bosonic ladder operator algebra with commutation relation [*a*_*i*_, *a*_*j*_] = 0, 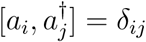 for the input population, allowing us to construct the input particle number eigenstates *|I*_*i*_⟩ obeying 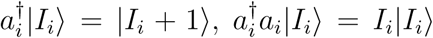. A Fock state with *I*_*i*_ particles on site *i* is obtained from the vacuum state *|*0⟩, defined by the relation *a |*0⟩ = 0, and 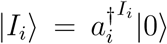. Similarly, we introduce annihilation and creation operators for output particles *b*_*i*_ and 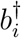 that commute with the input ladder operators: 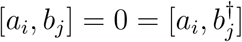.

Stochastic kinetics for the entire lattice is implemented by considering the master equation for the configurational probability *P* ({*I*_*i*_}, {*O*_*j*_}, *t*⟩, given by a sum over all lattice points on the right hand side of Eq.(9), by noting that a general Fock state is constructed by the tensor product *|*{*I*_*i*_}, {*O*_*j*_}⟩ = Π_*i*_*|I*_*i*_⟩|*O*_*i*_⟩. We define a time dependent formal state vector through a linear combination of all possible Fock states, weighted by their configurational probability at time *t*,

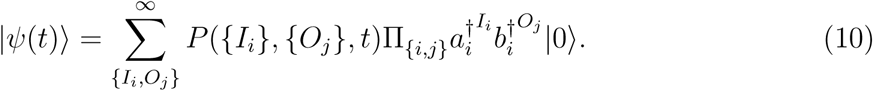

This superposition state encodes the stochastic temporal evolution. We use standard methods to transform the time dependence from the linear master equation into an imaginary time Schrödinger equation, governed by a time-dependent stochastic evolution operator *H*,

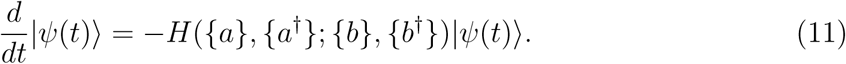

We may multiply Eq.(9) by 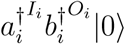, and sum over all values of *I*_*i*_, *O*_*i*_. With the definition of the state *|Ψ*(*t*)⟩,

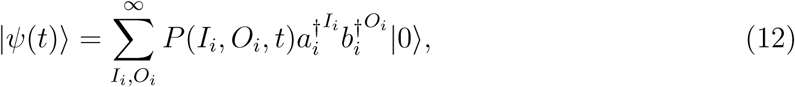

the *γ*_*I*_ term, i.e. *γ*_*I*_[(*I*_*i*_ + 1)*P* (*I*_*i*_ + 1, *O*_*i*_, *t*) − *I*_*i*_*P* (*I*_*i*_, *O*_*i*_, *t*)], in Eq.(9) becomes,

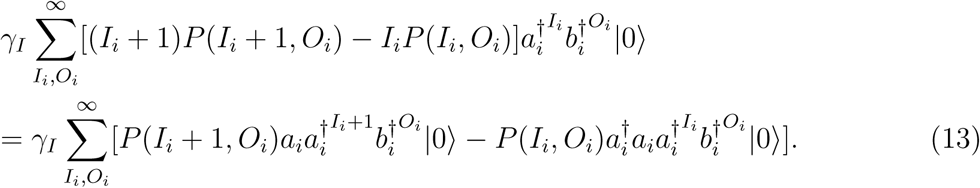

By relabeling the indices in the first sum, we arrive at the desired Hamiltonian expressed in second quantized representation as, 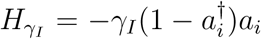. Similarly, terms with coefficients *F*, *γ*_*O*_ and *R*(*I*_*i*_) in Eq.(9) give the following contributions, 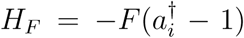, 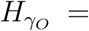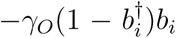, 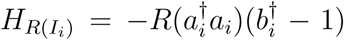. The total Hamiltonian *H* takes the following form,

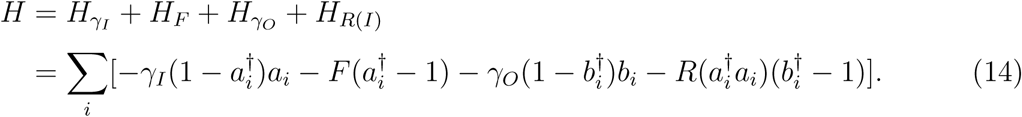

A convenient choice for the initial configuration for the master equation describing the stochastic particle reactions is an independent Poisson distribution at each site,

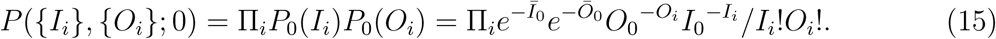

with mean initial input and output concentrations 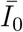 and 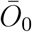. Just as in quantum mechanics, Eq.(11) can be formally solved leading to,

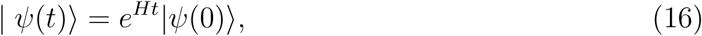

with the initial state 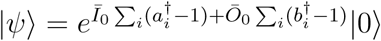.

Our goal is to compute averages and correlation functions with respect to the configurational probability *P* ({*I*_*i*_}, {*O*_*i*_}; *t*), which is accomplished by means of the projection state 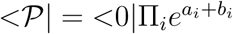, for which 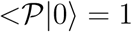 and 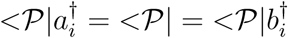, since 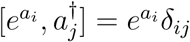. The average value of an observable *A*({*I*_*i*_}, {*O*_*i*_}) is,

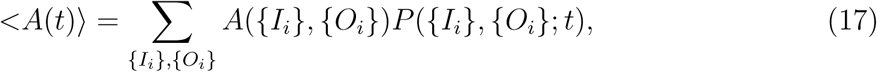

from which the statistical average of an observable can be calculated using,

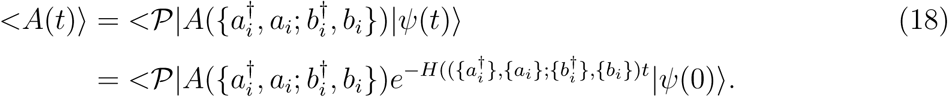

We follow a well-established route in quantum many particle theory [36], and proceed towards a field theory representation by constructing a path integral equivalent of the time dependent Schrödinger equation (Eq.(11)) based on coherent states [37]. These are defined as right eigenstates of the annihilation operators, *a*_*i*_*|α*_*i*_⟩ = *α*_*i*_*|α*_*i*_⟩ and *a*_*i*_*|β*_*i*_⟩ = *β*_*i*_*|β*_*i*_⟩, with complex eigenvalues *α*_*i*_ and *β*_*i*_. The coherent states satisfy 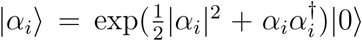, the overlap integral 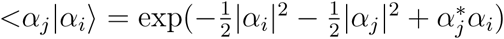, and the completeness relation ∫ Π_*i*_*d*^2^*α*_*i*_*|*{*α*_*i*_}*⟩<*{*α*_*i*_}*|* = *Π*. After splitting the temporal evolution (Eq.(11)) into infinitesimal increments, inserting the completeness relation at each time step, and with additional manipulations leads to an expression for the configurational average,

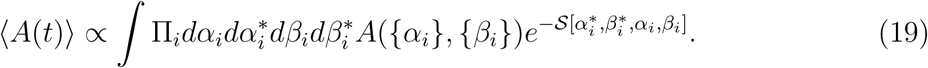

The exponential statistical weight is determined by the action,

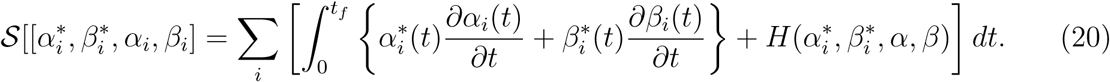

Finally, by taking the continuum limit using 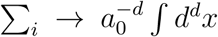, *a*_0_ is a lattice constant, *α*_*i*_(*t*) *→ ϕ*(*x, t*), *β*_*i*_(*t*) *→ ϕ*(*x, t*) and 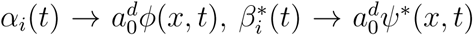, the expectation value is represented by a functional integral,

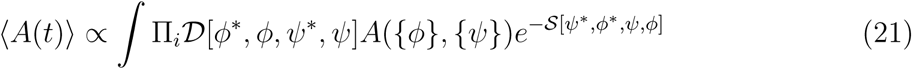

with an effective action

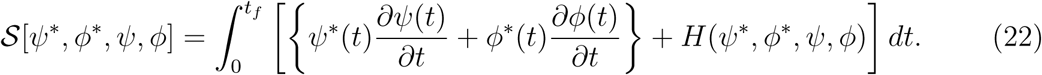

In the Hamiltonian (Eq.(14)), *a*^†^ and *b*^†^ are replaced by the field variables *ϕ*^∗^ and *ϕ*^∗^, respectively. Similarly, *a* and *b* operators become *ϕ* and *ψ* respectively.

The action in Eq.(22) encodes the stochastic master equation kinetics through four independent fields (*ψ*^∗^, *ϕ*^∗^, *Ψ, ϕ*). With this formulation, an immediate connection can be made to the response functional formulation using the Janssen − De Dominicis formalism for Langevin equations [38, 39]. In this approach, the response field enters at most quadratically in the pseudo-Hamiltonian, which may be interpreted as averaging over Gaussian white noise. With this in mind, we apply the non-linear Cole-Hopf transformation [40, 41], in order to obtain quadratic terms in auxiliary fields, 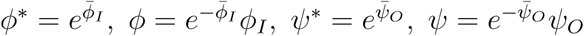, to the action in Eq.(22). The Jacobian for this variable transformation is unity, and the local particle densities are *ϕ*^∗^*ϕ* = *ϕ*_*I*_ and *ψ*^∗^*ψ* = *ψ*_*O*_. We obtain the following Hamiltonian,

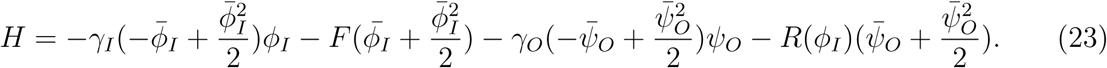

In the above equation, the exponential term has been expanded to second order. The rate equations are obtained through 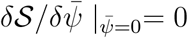 and 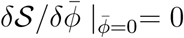. The terms quadratic in the auxiliary fields (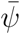 and 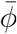) encapsulate the second moment of the Gaussian white noise with zero mean.

In order to obtain fluctuation corrections needed to calculate minimum error in signal transduction, we write the action in terms of fluctuating fields, *δϕ*_*I*_ = *ϕ*_*I*_ − 〈*ϕ*_*I*_〉 and *δΨ*_*O*_ = *ψ*_*O*_ − ⟨*ψ*_*O*_⟩ as,

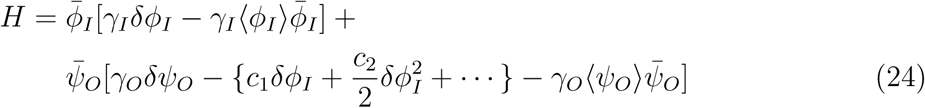

where we have expanded *R*(*ϕ*_*I*_) in a Taylor series,

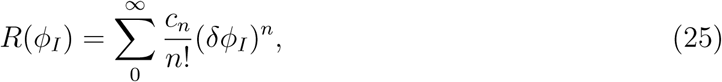

with constant *c*_*n*_. Note this expansion differs from the one used in Eq.(6). The coefficients of 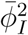 and 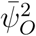 reflect the noise correlations in Langevin description.

In Fourier space the action becomes

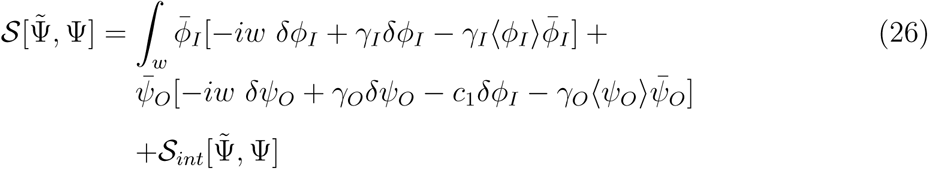

where 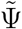 represents the set 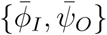 and Ψ denotes {*ϕ*_*I*_, *ψ*_*O*_}. The non-linear contribution to the action is 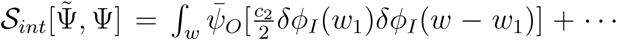. Physical quantities can be expressed in terms of correlation functions of fields Ψ and 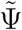, taken with the statistical weight 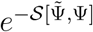,

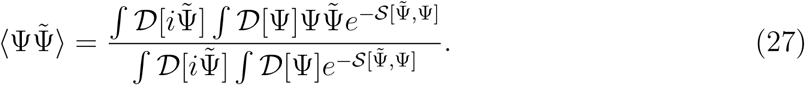

In order to compute the correlation function involving response fields, it is useful to introduce the generating functional,

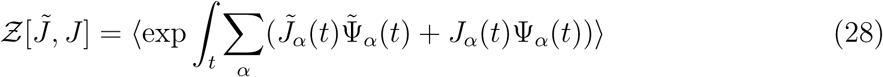

where *α* represents the set {*ϕ*_*I*_, *ψ*_*O*_}, for which the required correlation functions are obtained via functional derivatives of *Ƶ* with respect to the appropriate source fields.

The procedure is readily implemented for the Gaussian theory with statistical weight 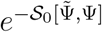. In Fourier space, we can write the harmonic function as,

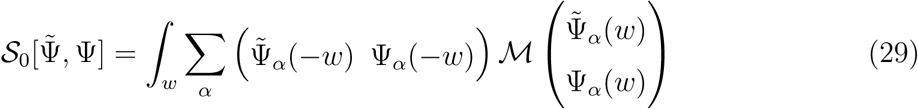

with the Hermitian coupling, a (4,4) matrix *ℳ*(*w*). With the aid of Gaussian integrals, we obtain,

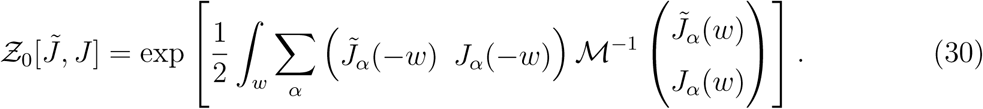

From Eq.(30), we now directly infer the matrix of two point correlation functions in the Gaussian ensemble with the inverse of harmonic coupling matrix *ℳ*.

## APPLICATIONS

As a first application we apply the field-theoretic formalism to the push-pull network, which can be exactly solved for the error (Eq.(5)). In the process we illustrate the way the diagrammatic expansion works in the context of signaling networks, making it possible to apply the theory to more complicated systems.

### A. Push-Pull network

The calculation of the error (Eq.(5)) in terms of the control variable (the average number of phosphatase molecules per cell 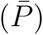) requires the correlation functions ⟨*δOδI*⟩, ⟨*δO*^2^⟩ and ⟨*δI*^2^⟩. These can be expressed in terms of the matrix elements of (*ℳ*^−1^)_*mn*_ (Eq.(29)). Subscripts *ℳ* and *n* represent the *ℳ*^*th*^ row and *n*^*th*^ column, respectively. For example, (*ℳ*^−1^)_33_ is the correlation function ⟨*δϕ*_*I*_(*−w*)*δϕ*_*I*_(*w*)⟩. Similarly we can obtain other correlation functions. Now we can compute, power spectra for the input and output molecules by evaluating the correlation functions of kinase and substrate populations by using Eq. (30). We use perturbation theory for the action corresponding to the push-pull network to compute the non-linear contribution to the correlation function.

We obtain the following expressions for the power spectra,

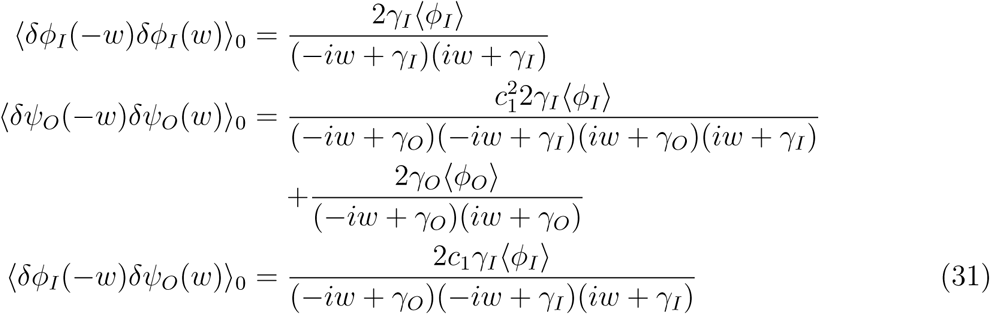

The ⟨…⟩_0_ is taken with respect to the non-interacting theory 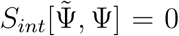 in Eq.(26)). Using these functions, the error (*E*) and gain (*G*) are given by,

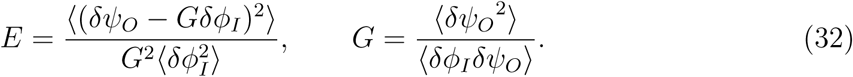

By inserting the expressions for the correlation functions in Eq.(31) into Eq.(32), and integrating over *w*, we obtain the minimum relative error for the linear push-pull network,

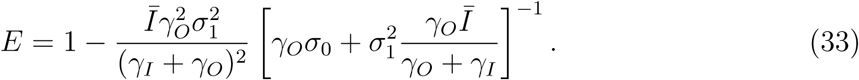

Higher order corrections to the power spectra ⟨*δΨ*_*O*_(−*w*)*δΨ*_*O*_(*w*)⟩ are calculated using perturbation theory by evaluating the Feynman diagrams (Fig.(2)),

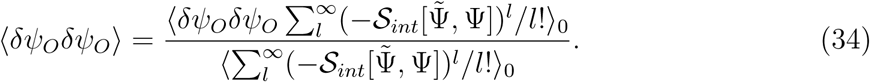

**FIG. 2:**
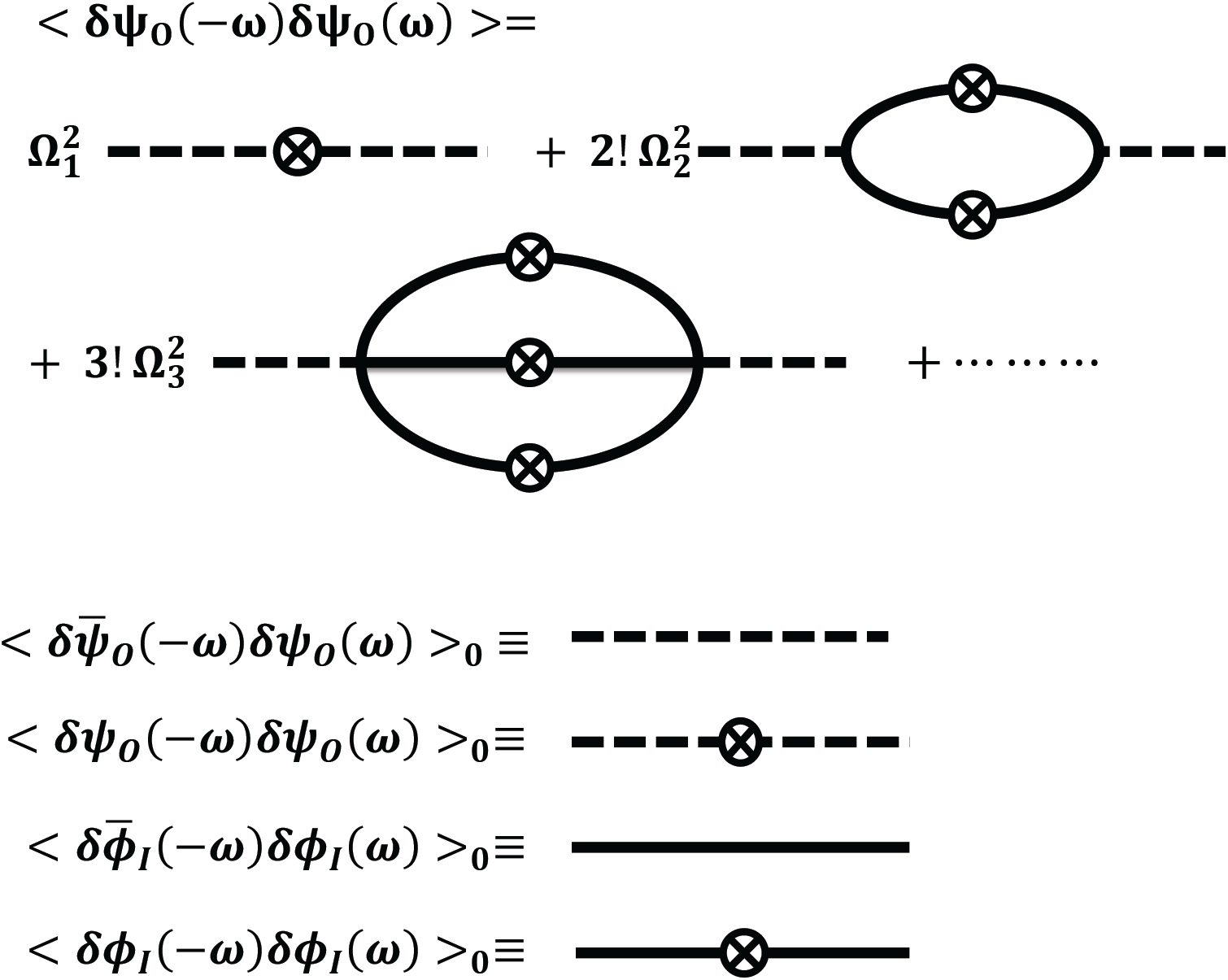
Examples of diagrams for the correlation function ⟨*δΨ*_*O*_(*−w*)*δΨ*_*O*_(*w*)⟩. The Ω_*n*_s are coefficients with, 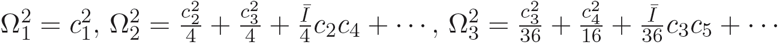 and so on.

For example, the second order contribution to the ⟨*δΨ*_*O*_(−*w*)*δΨ*_*O*_(*w*)⟩ arising from the loop in Fig.(2) is 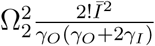 (see Appendix A for details). The coefficient 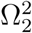 is given by 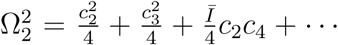. Higher order terms have a similar structure: for example, the third order contribution to the power spectra is 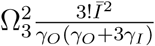, with 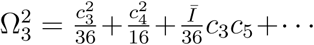. By evaluating all the diagrams in Fig.(2), we obtain the final expression for the relative error,

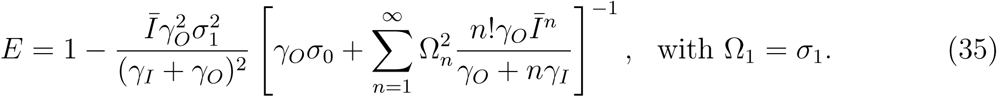

The form of the result in Eq.(35) coincides with the exact expression (Eq.(7)) for the relative error previously obtained [18] by using an entirely different approach based on umbral calculus. However, the coefficients Ω_*n*_ are expressed in terms of the coefficients *c*_*n*_ used in the series for *R*(*I*) (Eq.(25)) rather than *σ*_*n*_. The two kinds of coefficients are non-trivially related through,

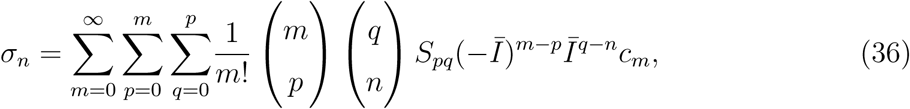

where *S*_*pq*_ are Stirling’s numbers of second kind. For all *n*, the leading order term 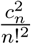 of Ω_*n*_ is the same as the leading order term of *σ*_*n*_.

The sum within the bracket in Eq.(35) is composed of non-negative terms. The minimal sum *E* is obtained by setting Ω_*n*_ = 0 for all *n ≥* 2. Thus, *E* is bounded from below by 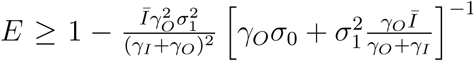. The term on the right hand side is minimized with respect to *γ*_*o*_ when 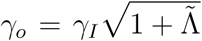, with 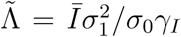. At the optimal *γ*_*O*_, the equality becomes 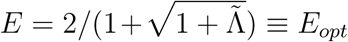. As *σ*_1_ increases, 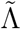 becomes large which is desirable for high fidelity signal transduction. As long as *R*(*I*) is approximately linear in the vicinity of 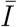, the corrections *σ*_*n*_ (or Ω_*n*_) for *n >* 2 are negligible, and *E* is close to *E*_*opt*_. The coefficients *σ*_*n*_ for *n >* 2 must be non-negligible when *σ*_1_ is sufficiently large. Such a highly sigmoidal input-output response, known as ultra-sensitivity [1], is biologically realizable in certain regimes of signaling cascades. In the limit of a nearly step-like response, non-linearity in *R*(*I*) becomes appreciable around 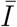, distorting the output signal and leading to *E* that is larger than *E*_*opt*_. Because *E* increases with 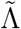 in this limit, the benefits of ultra-sensitivity vanish.

### B. Signaling Cascades

A natural extension is to consider a cascade created by an array of connected push-pull networks. Indeed, in some biological signaling pathways external perturbation is transmitted through a cascade of reactions involving successive activation by kinases and deactivation by phosphatases. An example is the stimulation of a receptor tyrosin kinase by epidermal growth factor, which results in downstream responses of the MAPK network [27, 42].

Because sections *B*, *C* and *D* are related, we explain briefly the results in order to ensure that the relationship between these sections are clear. In this section we describe the two cascade network using the field theory framework, and the coarse-graining procedure needed for obtaining an analytic expression for optimal error. In section *C*, we show that the two cascade network behaves as noise filter with a time delay, *α*^−1^. By mapping the cascade to a push-pull network with an intermediate, we show in section *D* that *α* can be exactly calculated. Thus, the results in the three sections provide an analytic theory for optimal signaling in the two cascade network.

Consider a two step series enzymatic cascade (Fig.(3)) modeled as a sequence of two enzymatic push-pull loops stimulated by an upstream enzyme. In the first loop, an upstream enzyme, *K* phosphorylates the substrate, *S*, to produce *S^∗^*, converting it from an inactive to active state. Phosphatase (*P*) dephosphorylates *S*^∗^ to an inactive state *S*. In the second loop, *S*^∗^ acts as the enzyme for the phosphorylation of *T* and *P*, the corresponding phosphatases. The series of chemical reactions involved in this cascade are,

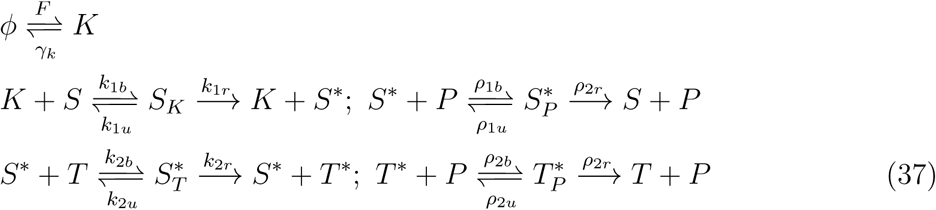

where *S*_*K*_, 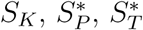, and 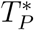 are the reaction intermediates, and *k*_*ib*_, *k*_*i*_*u*, *k*_*ir*_, *ρ*_*ib*_, *ρ*_*i*_*u* and *ρ*_*ir*_, *i* = 1, 2, are the rate constants of the stochastic biochemical reactions in the cascade. The input signal *K* + *S*_*K*_ is transduced into the active substrate output 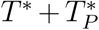. In an insightful article [42], a deterministic approach was used to analyze the system of chemical reactions in Eq.(37). Here we assume that the reactions are stochastic. In order to develop analytical results we only consider fluctuations of all species that deviate linearly from their mean values. The validity of the asumption is established by comparing the results with kinetic Monte Carlo (KMC) simulations.

**FIG. 3:**
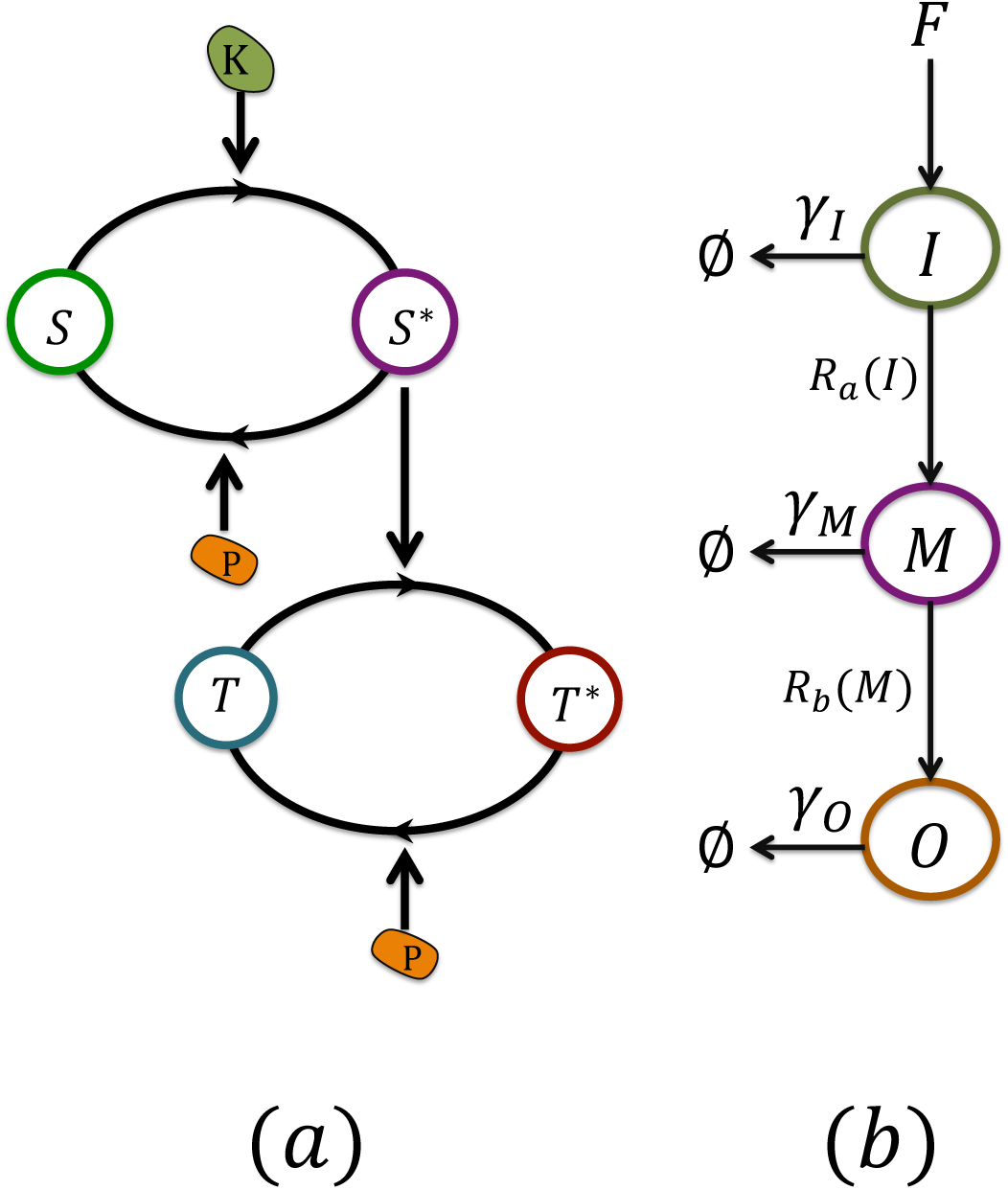
(a) Enzymatic cascade that arises naturally in mitogen activated protein kinase (MAPK) networks. In a caricature of such a network, kinase (*K*) phosphorylates the substrate (*S*), leading to the formation of *S^∗^*. Deactivation is triggered by reactions with the phosphatase (*P*). *S*^∗^ phosphorylates the substrate *T*, producing *T*^∗^ and *P* reverts it to the original form through dephos-phorylation. The rate parameters in the chemical reactions (Eq. (37)) used to produce numerical results (in units of *s*^−1^) are : *k*_1*b*_ = *k*_2*b*_ = *ρ*_1*b*_ = *ρ*_2*b*_ = 10^−5^, *k*_1*u*_ = 0.02, *k*_2*u*_ = 0.3, *ρ*_1*u*_ = 0.5, *ρ*_2*u*_ = 1.0, *k*_1*r*_ = 3, *k*_2*r*_ = 5.0, *ρ*_1*r*_ = 0.3, *ρ*_2*r*_ = 0.1, *F* = 1. The deactivation rate *γ*_*k*_ = 0.01*s*^−1^ controlling the characteristic time scale over which the input signal varies. Mean free substrate and phosphatase populations are in the ranges 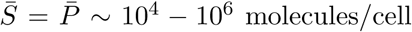. (b) Three species coarse-grained signaling network with the indicated rates is intended to capture the physics of the cascade in (a). The mathematical equivalence between the networks in (a) and (b) is established in the text.

For the network in Fig.(3), the procedure outlined earlier leads to a Schrödinger-like equation with the following Hamiltonian,

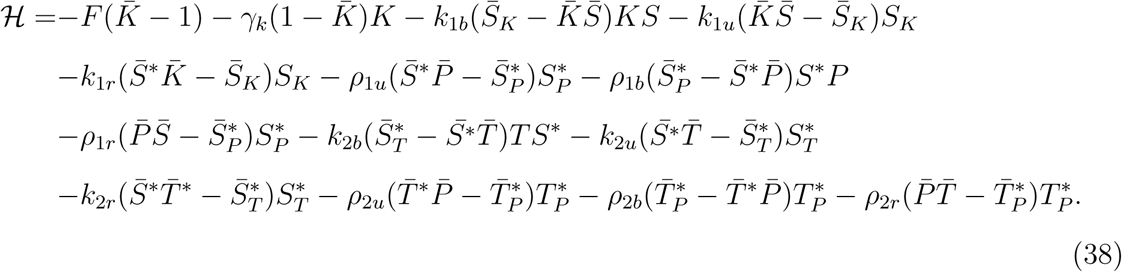

We can approximately map the two-step cascade into a two-species coarse-grained network, which acts like a noise filter, as described in detail in Ref. [18]. Consider a signaling pathway (Fig.(1)) with time varying input *I*(*t*) and time varying output *O*(*t*). These are the total populations (free and bound) of the input and output active kinases, with *I* = *K* + *S*_*K*_ and 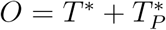. The upstream pathway provides an effective production rate *F* of input *I*, while the output *O* results from the reaction 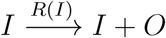. As before, *γ*_*I*_ and *γ*_*O*_ are the degradation rates for the input and output respectively, mimicking the role of phosphatase. The input and output correlation functions, evaluated using the field theory formalism, have the approximate structure,

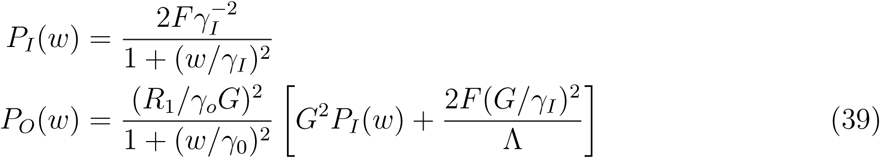

where we have used a linear approximation for 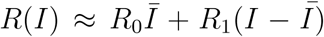 with *R*_0_, *R*_1_⟩0. Optimality is achieved when 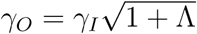 with gain 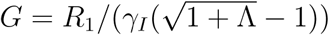. Relative error with the minimum 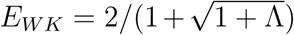. As before, the fidelity between output and input is controlled by single dimensionless control parameter Λ = (*R*_1_*/γ*_*I*_)(*R*_1_*/R*_0_)^2^. This mapping allows us to use the general WK result for gain (*G*) and the minimum relative error (*E*_*W*__*K*_) to predict the optimality condition, allowing us to calculate the minimum possible value of *E*. The results for the error in terms of the mean number of phosphatase are given by the red lines in Fig.(4).

**FIG. 4:**
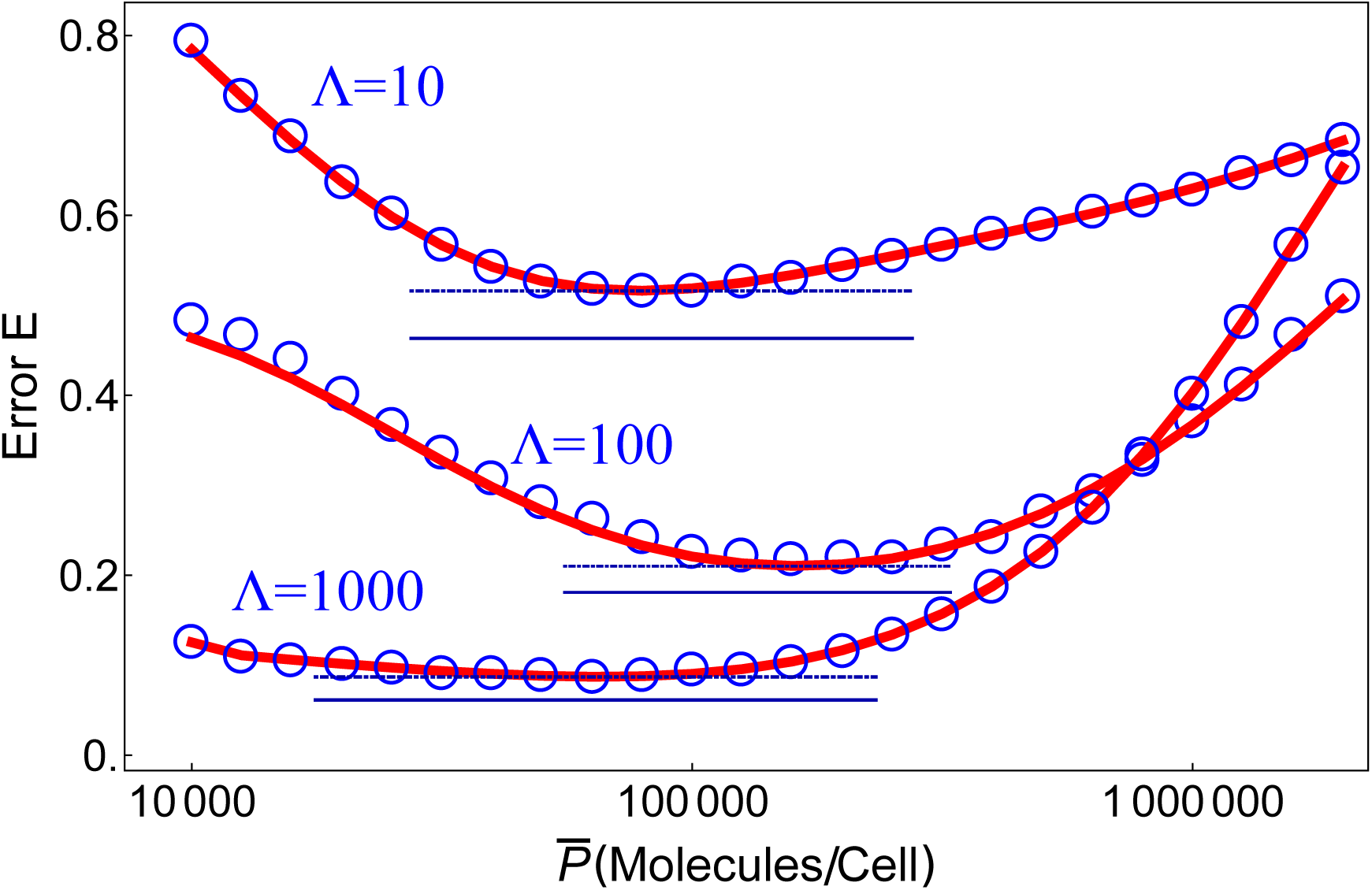
Relative error *E* for the signaling cascade (red lines show the theoretical predictons; blue circles are obtained using the kinetic Monte Carlo simulations) for three λ values. The blue dashed line gives the predictions (Eq.(44)) using the WK formalism for *E*_*W*__*K*_ with time delay. The solid blue line is the minimal error corresponding to the theory without time delay in Eq.(5). The comparison shows that the two-loop cascade behaves as a push-pull network with a time delay. The time delay parameter, *α*, is explicitly given in Eq.(47). Thus, the theory has no adjustable parameter.

In order to test the accuracy of our theory we simulated the dynamics of the enzymatic cascade using the KMC method. The relative error *E* shown in Fig.(4) is in excellent agreement with the theoretical predictions. Interestingly, *E* achieves a minimum at 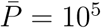 molecules/cell, which is ten times larger than the phosphatase concentration in the one step enzymatic push-pull loop using similar parameters. Fig.(4) shows that there is a well defined narrow range of phosphatase population in which the error is minimum. The range decreases as λ decreases (Fig.(4)). The minimum value for the relative error does not reach the value predicted by the WK limit (Eq.(5)). As we show below, the additional error arises from an effective time delay as the signal passes from one cascade to another. We also demonstrate that the time delay can alternatively be mimicked by reducing the two cascade system to a coarse-grained pathway with an intermediate (Fig.(3b)).

**FIG. 5:**
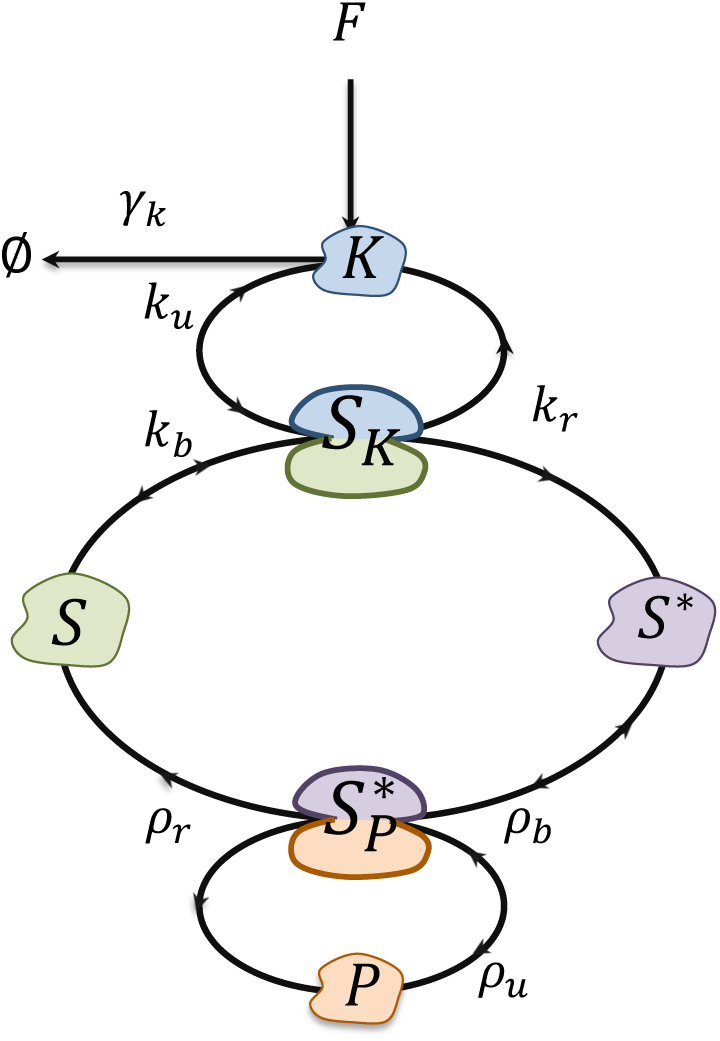
Enzymatic push-pull loop showing phosphorylation of the substrate (*S*) by kinase (*K*) to produce the active form *S*^∗^. Phosphatase (*P*) reverts it to the original form through dephosphorylation. *S*_*K*_ and 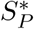 represent the substrate in the complex with kinase and phosphatase, respectively. Binding, unbinding and the reaction rate constants are shown with arrows.

### C. Noise filtering with time delay

In order to prove that the two-cascade loop effectively acts like a noise filter with time delay, we derive the condition for minimum error for the latter following the Bode-Shannon formulation of the WK theory [11]. In this scenario, the transmitted signal can only be recovered after a constant delay, *α*. The output *O*(*t*) is produced by convolving the corrupted signal (input *GI*(*t*)+ noise *n*(*t*)) with a causal filter *H*(*t*). In Fourier space, we obtain,

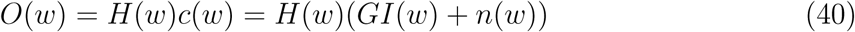

where 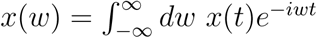 for the time series *x*(*t*). The relative error is given by [11],

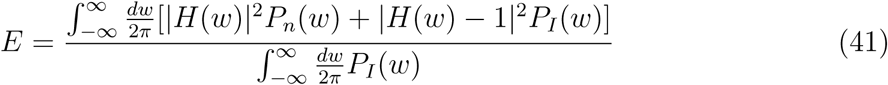

where *P*_*I*_(*w*) and *P*_*n*_(*w*) are the power spectral densities (PSDs) of *GI*(*t*) and *n*(*t*) respectively. We need to minimize *E* in Eq.(41) over all possible *H*(*w*), with the condition that *H*(*t*) = 0 for *t < α*. The optimal causal filter has the following form [9, 11, 22],

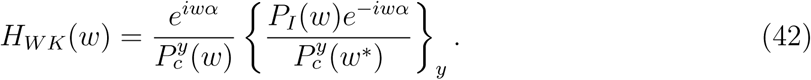

The *y* super and subscript refer to two different decompositions in the frequency domain. Causality can be enforced by noting the following conditions: (i) Any physical PSD, in this case *P*_*c*_(*w*) corresponding to the corrupted signal *c*(*t*) = *GI*(*t*) + *n*(*t*), can be written as 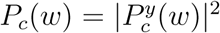. The factor 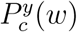, if treated as a function in the complex *w* plane, does not have zeros and poles in the upper half-plane (Im *w*⟩0). (ii) We also define an additive decomposition denoted by {*F* (*w*)}_*y*_ for any function *F* (*w*), which consists of all terms in the partial fraction expansion of *F* (*w*) with no poles in the upper half-plane. By using the PSDs, 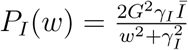 and 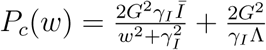, we obtain the following optimal filter *H*_*W K*_ (*w*),

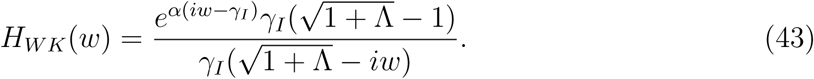

In the limit 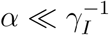, the optimal error *E*_*W*__*K*_ takes the following form [22],

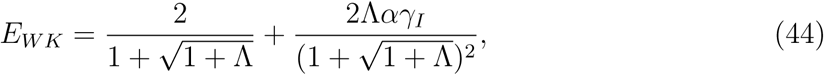

where second term in the above equation is the correction due to the time delay to the WK minimum value of the relative error for an instantaneous filter (*α →* 0). The correction is positive for all values of *α* and λ, which implies that time delay must increase the error in signal transmission. If we add this correction to the WK minimum result for the relative error of instantaneous filter (Eq.(5)), for specific values of *α* calculated explicitly in the following section, we recover the minimum relative error in the signaling cascade. Thus, the two step enzymatic cascade minimizes the noise but behaves like a single step network with a time delayed filter.

### D. Deriving the time delay *α* by mapping onto a three-species pathway with an intermediate

Alternatively, we can derive an explicit expression for the delay parameter *α* by using a different mapping for the original cascade. Instead of mapping onto a twospecies network of *I* and *O* with a time delay, we map onto a three-species network (Fig.(3b)) with *I*, *ℳ*, and *O*. Here there is no explicit time delay, but an additional species *ℳ* that will play the role of a “pseudo” intermediate mimicking the effect of the time delay. This network is governed by the reactions: 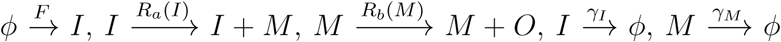 and 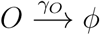. The production functions have the linear form: 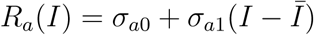 and 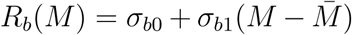. Earlier analysis of this network [22] has shown that it behaves like a time delayed filter, with the minimal error in the same form as Eq.(44), with 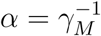 and effective 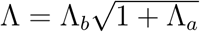, where 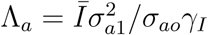 and 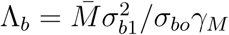.

The original signaling cascade (Fig.(3a)) can be mapped onto the three-species pathway (Fig.(3b)). This involves identifying the population 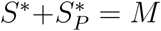 as a “pseudo” intermediate, with an effective degradation *γ*_*ℳ*_. The mapping can be carried out by comparing PSDs between the two models. For the three-species network these are given by,

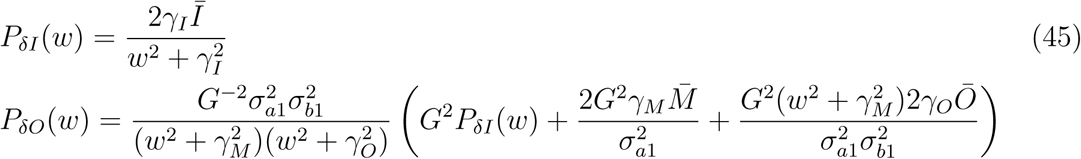

Now, the PSDs for signaling cascade calculated from Doi-Peliti formalism are given by

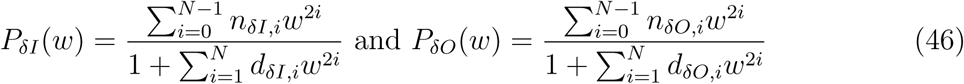

where the *w*-independent parameters *n*_*δI,i*_, *n*_*δO,i*_, *d*_*δI,i*_ and *d*_*δO,i*_ are related to the rate coefficients in the cascade reactions (Eq.(37)). Here, *N* = 7 corresponds to the number of independent dynamical variables 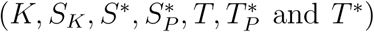. By mapping Eq.(46) into Eq.(45), we can extract the degradation rate of intermediate species 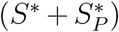, *γ*_*ℳ*_ in terms of coefficients in Eq.(46),

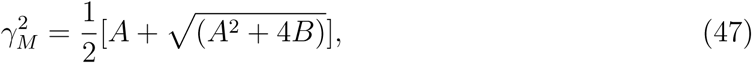

with 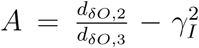 and 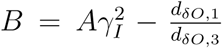. The time delay parameter 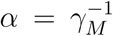 in the signaling cascade. With this identification for *α* we have a complete theory for *E*, with no adjustable parameter, as a function of the control parameter, the mean phosphatase levels. It is tempting to speculate that a multiple (*>* 2) step cascade might also be mathematically equivalent to a network with a single pseudo intermediate.

### E. Enzymatic Push-Pull Loop

In considering the cascade model, we focused on the case where fluctuations around mean populations levels were small enough that the linear approximation is valid. To study the effects of non-linearity, we will look at a simpler system (one stage of the cascade) but without any constraints on the size of the fluctuations. A microscopic model for the enzymatic push-pull network is shown in Fig.(5). The upstream enzyme, *K* phosphorylates a substrate *S* to *S*^∗^, thereby converting it from an inactive to an active state. The effective production rate in the upstream pathway for enzyme *K* is *F*. The degradation rate for *K* is *γ*_*K*_. The enzyme is either free (*K*) or bound to substrate (*S*_*K*_). The input *I* is the total enzyme population *I* = *K* + *S*_*K*_. Phosphatase, *P*, on the other hand dephosphorylates the active substrate *S*^∗^ to an inactive state *S*. The output of the two phosphorylation cycle is 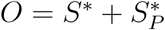.

The biochemical reactions for the enzymatic network with the corresponding rate constants are,

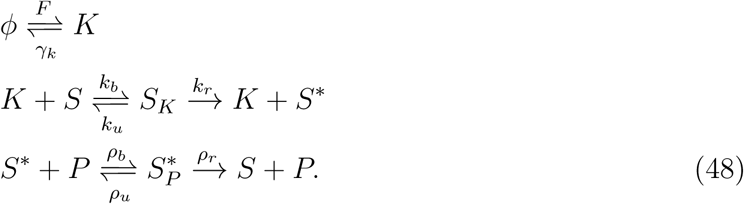

In the stochastic chemical reactions that govern the phosphorylation/dephosphorylation steps, the input signal *I* = *K* + *S*_*K*_ is transduced into the active substrate output 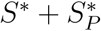. To derive the conditions for optimality, we follow the procedure outlined in the previous section. Starting from the master equation, we can derive a Schr*ö*dinger-like equation with the following Hamiltonian,

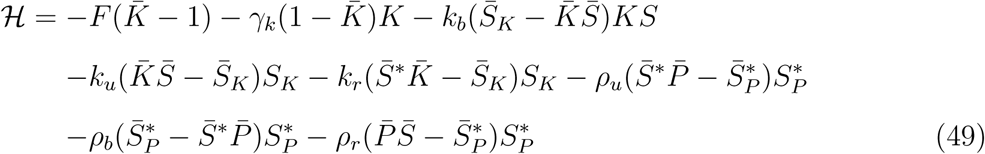

The field variables 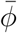 are associated with creation operators of corresponding population. Similarly *ϕ* correspond to annihilation operators. After using coherent-state path integral formalism, we arrive at the expression for the action corresponding to the enzymatic push-pull loop from which we calculate the power spectra for the input and output.

As in the signaling cascade network described in the previous section, we approximately map the complete enzymatic network into a noise filter [18]. The input and output correlation functions, evaluated using field theory formalism, have the approximate structure given in Eq.(39). Starting from the full dynamical equations (Eq.(48)), we compute correlation functions using field theory by solving the Wiener-Hopf relation in Eq.(3), for the optimal function *H*_*W*__*K*_(*t*).

Correlation functions of input and output calculated for enzymatic push-pull loop have the approximate form of Eq.(39), with effective values of parameters *γ*_*I*_, *γ*_*O*_, *R*_1_ and Λ which have been expressed in terms of loop reaction rate parameters. This mapping allows us to use WK result for the gain (*G*) and minimum relative error (*E*_*W*__*K*_) to predict the optimality and minimum possible value of *E*. The results for the error in terms of the mean number of phosphatase are given by the solid lines in Fig.(6).

**FIG. 6:**
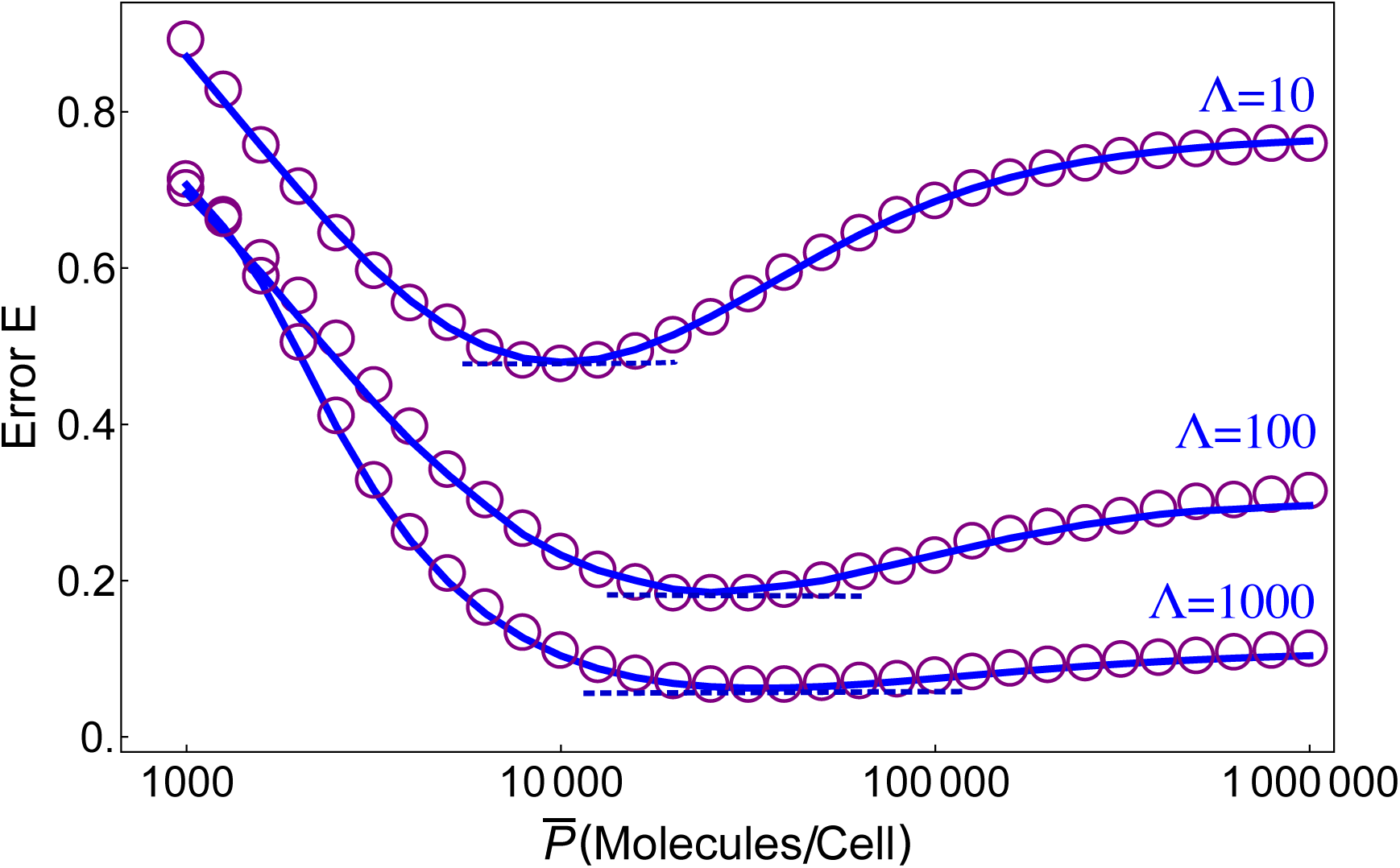
Relative error *E* for the enzymatic push-pull loop in Fig.(5). The blue lines correspond to theoretical predictions. The KMC simulation results are given in purple circles. The dashed line is the minimal error corresponding to the WK theory (Eq.(5)). The values of the rates corresponding to the chemical reactions in the enzymatic push-pull network (Eq.(48)) is given in the main text. For the parameter values the predictions of the linear theory are very accurate.

In order to illustrate the accuracy of the theory we performed KMC simulations by choosing the forward and backward reaction rates in Eq.(48) describing the enzymatic push-pull loop network (all units are in *s*^−1^) : *k*_*b*_ = *ρ*_*b*_ = 10^−5^, *k*_*u*_ = 0.02, *ρ*_*u*_ = 0.5, *k*_*r*_ = 3, *ρ*_*r*_ = 0.3, *F* = 1. The deactivation rate *γ*_*k*_ = 0.01*s*^−1^ of enzyme *K* which controls the characteristic time scale over which the input signal varies, mimicking the role of phosphatase. Mean free substrate and phosphatase populations are in the ranges 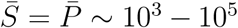 molecules/cell. Fig (6) shows that *E* is a minimum at a particular value of phosphatase concentration 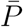, where optimality condition is satisfied i.e. 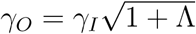. For a particular value of λ = 100, we see minimum error *E* = 0.18 for the enzymatic push-pull loop. The result of the KMC simulations (purple circles) are in excellent agreement with the analytical calculation (blue line) for all Λ values.

In the parameter space used in the results in Fig.(6), a linear theory reproduces the simulation results well. However, deviations from the predictions of the linear theory are expected if the input parameters are varied. In order to investigate these deviations we first obtained the error using the parameter values, *k*_*b*_ = *ρ*_*b*_ = 10^−3^, *k*_*u*_ = 0.02, *ρ*_*u*_ = 0.5, *k*_*r*_ = 3, *ρ*_*r*_ = 0.3 using KMC simulations. The relative error for λ = 100 is shown in purple line in Fig.(7). The blue line, calculated from linear theory predictions, deviates substantially from simulations (purple line in Fig.(7)). To improve the predictions of the theory we calculated second order corrections to *E*. The result, displayed as green curve in Fig.(7)), shows that there is improved agreement between theory and simulations. The non-linear corrections, which are substantial, brings the theoretical predictions closer to the simulation results, especially near the values of 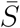 for which the error is a minimum (Fig.(7))). We suspect that higher order perturbative corrections will further improve the results based on the following observation. We fit the dependance of the error for 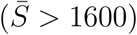 using the function, 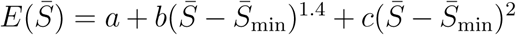, where *a, b* and *c* are constants and 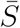 is the value of 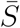 at which 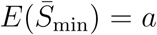 is a minimum. The functional form of 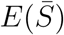 is the same for the exact simulation results, and the predictions of the linear and non-linear theory except the coefficients *a, b* and *c* are different. We, therefore, surmise that higher order terms merely renormalize the coefficients, keeping unaltered the form of relative error. Consequently, we conclude that improved estimates of *a, b*, *c* from third and higher order contributions should produce predictions in better agreement with simulation.

**FIG. 7:**
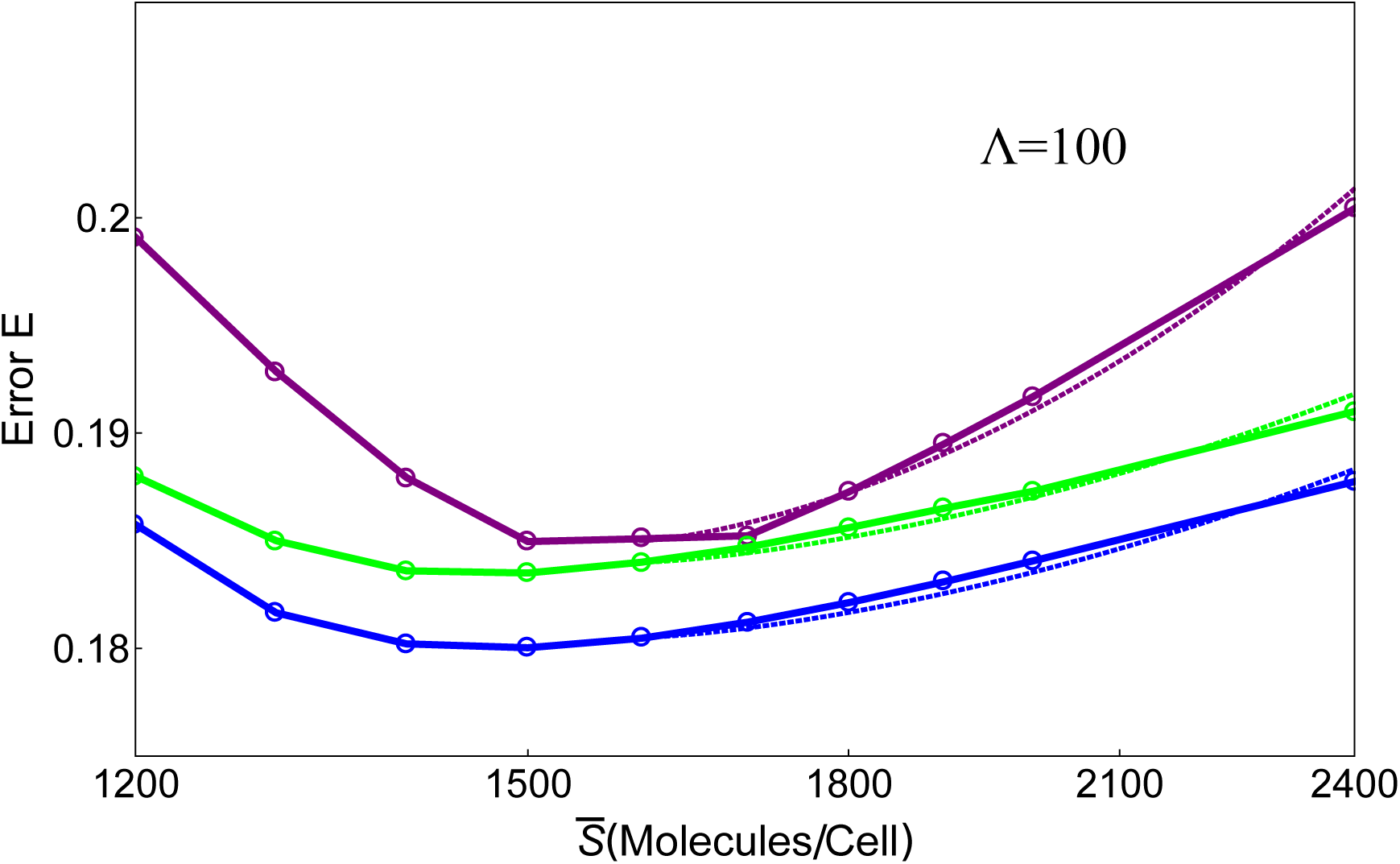
Error *E* for the enzymatic push-pull loop for different values of 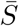 with λ = 100. The rate parameters used in Eq.(48) are *k*_*b*_ = *ρ*_*b*_ = 10^−3^, *k*_*u*_ = 0.02, *ρ*_*u*_ = 0.5, *k*_*r*_ = 3 and *ρ*_*r*_ = 0.3. The blue line is the result calculated using linear theory. The green line results from second order corrections to the error *E*. The KMC simulation results are given in purple line. Clearly inclusion of non-linear corrections improves the predictions of the theory in the range of 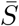 values for which *E* is small. Dotted lines are fit with the function 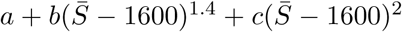 where *a*, *b*, *c* are constants. For all the curves *a*, *b*, *c* values change but the functional form of *E* as a function of 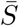 is the same. It is likely that if the theory is extended beyond second order, there should be further improvement by bringing *a*, *b* and *c* values closer to the simulation results.

## CONCLUDING REMARKS

In order to assess the accuracy of signal transmission, using the mean square of the error between input and output as a fidelity measure, we have developed a field theoretic formulation that allows us to predict conditions for optimal information transfer for an arbitrary stochastic chemical reaction network. The starting point is the classical master equation for interacting particle systems, which is mapped to a non-Hermitian ’quantum’ many-body Hamiltonian dynamics. Finally, the coherent-state path integral representation is utilized to arrive at a continuum field theory description that faithfully incorporates the intrinsic reaction noise and discreteness of the original stochastic processes. The formulation allows us to use standard field theory methods to compute the relative error in the information transfer using perturbation theory to all orders in non-linearity. This approach leads to an analytical expression for the minimum relative error in signal transduction. The usefulness of the general field theory formulation is illustrated through signaling networks of increasing complexity.

Detailed study of an enzymatic push pull loop, the basic unit involved in complex signaling pathways, show that it behaves like an optimal linear WK noise filter, as previously established using entirely different methods [18]. In this particular case, the joint probability *P* (*δI, δO*) is approximately bivariate Gaussian, which means the error *E* is also directly related to the mutual information *M* in bits between *δI* and *δO* as *E* = 2^−2*M*^ [22].

The two-stage enzymatic cascade behaves as an optimal filter without achieving the minimum predicted by the WK theory. We attribute the deviation to the time delayed response of the cascade. By mapping the cascade signaling network to a three-species push-pull like model with a pseudo intermediate state we derived an explicit expression for the time delay. We show that the time delay is associated with the degradation rate of the pseudo intermediate state in the coarse-grained representation of the two-step cascade. We also demonstrate that in those cases where the linear approximation breaks down, systematic perturbative corrections can be calculated using our theory, which minimize the difference between the findings in the simulations and theoretical predictions. The success in this example illustrates the power of the formalism. Analyzing experimental data using the framework introduced here will help decipher the design principles governing signaling networks in biology, and allow us to understand the constraints imposed by noise in information transfer.

## Acknowledgements

We are grateful to the National Science Foundation (CHE 16-61946) for supporting our work. Much of this work was carried out while the authors were in the Institute for Physical Sciences and Technology in the University of Maryland, College Park.

## Appendix A Second order loop correction to the signaling error for the push-pull network

Here, we illustrate the calculation of *E* arising from perturbation expansion of the field theory for the push-pull network with non-linearity explained in the text. To second order the diagram needed to compute *E* is,

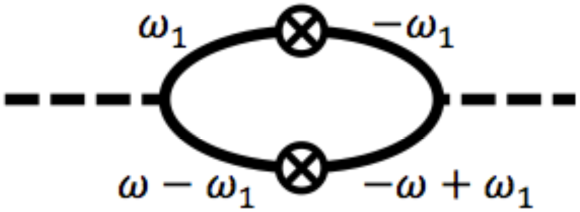

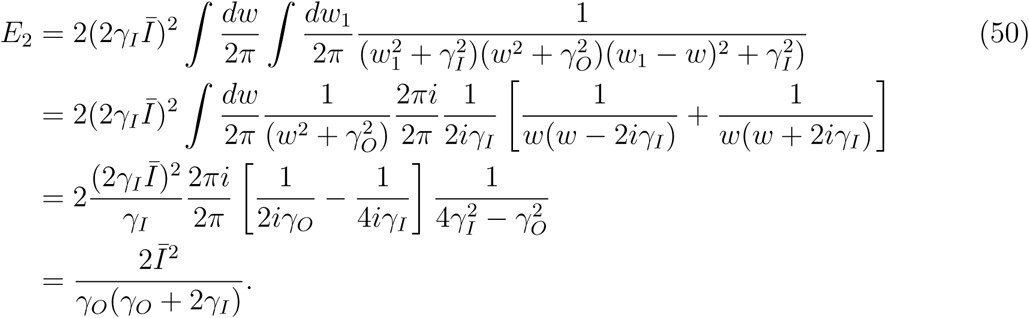

In the first line of the above equation, we perform the complex integration in the upper half plane by evaluating the residues at poles *w*_1_ = *iγ*_*I*_ and *w*_1_ = *w* + *iγ*_*I*_, respectively. Similarly, in the second line we calculate the residues at poles *w* = *iγ*_*O*_ and *w* = 2*iγ*_*I*_.

The coefficient, 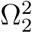 (Eq.(35)), is diagrammatically represented as,

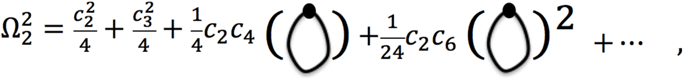

where the expression for the loop in the first bracket is 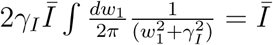. The coefficients Ω_*n*_ in Eq.(35) are functions of *c*_*n*_. In turn, Ω_*n*_s and *σ*_*n*_s are also connected by the relation between *σ*_*n*_ and *c*_*n*_ (see main text). For all *n*, the leading order term 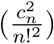 of Ω_*n*_ and *σ*_*n*_ is identical.

## Appendix B Action for enzymatic push-pull network

We give the form of the action here for the enzymatic push-pull network for which the chemical reaction scheme is given in Eq.(48). Despite the complexity, the action can be manipulated using Mathematica in order to obtain general expression for the error.

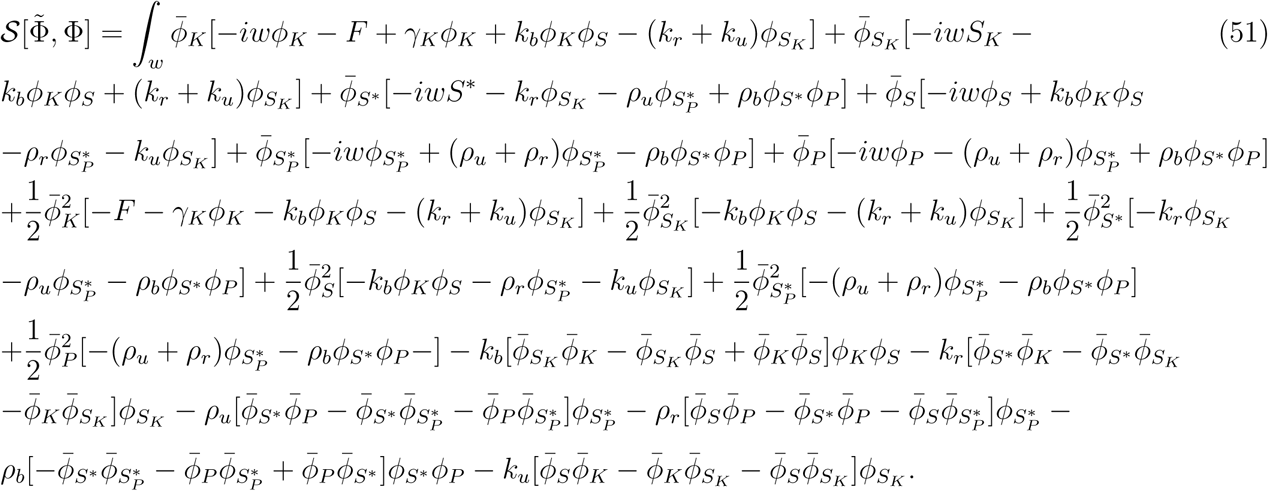

